# Multi-trait selection to build resilience in conifer forests: a case study on spruce-shoot weevil interactions

**DOI:** 10.1101/2022.01.21.477209

**Authors:** Jaroslav Klápště, Barry Jaquish, Ilga Porth

## Abstract

Tree planting programs now need to consider climate change increasingly, therefore, the resistance to pests plays an essential role in enabling tree adaptation to new ranges through tree population movement. The weevil *Pissodes strobi* (Peck) is a major pest of spruces and substantially reduces lumber quality. We revisited a large Interior spruce provenance/progeny trial (2,964 genotypes, 42 families) of varying susceptibility, established in British Columbia. We employed multivariate mixed linear models to estimate covariances between, and genetic control of, juvenile height growth and resistance traits. We performed linear regressions and ordinal logistic regressions to test for impact of parental origin on growth and susceptibility to the pest, respectively. A significant environmental component affected the correlations between resistance and height, with outcomes dependent on families. Parents sourced from above 950 m a.s.l. elevation negatively influenced host resistance to attacks, probably due to higher *P. engelmannii* proportion. For the genetic contribution of parents sourced from above 1,200 m a.s.l., however, we found less attack severity, probably due to a marked mismatch in phenologies. This clearly highlights that hybrid status might be a good predictor for weevil attacks and delineates the boundaries of successful spruce population movement. Families resulting from crossing susceptible with resistant parents generally showed fast-growing trees were the least affected by weevil attacks. Such results indicate that these “hybrids” might be genetically better equipped with an optimized resource allocation between defence and growth and might provide the solution for concurrent improvement in resistance against weevil attacks, whilst maintaining tree productivity.

## Introduction

Climate change, mainly caused by global warming through carbon dioxide emissions (greenhouse gas effect; [1]), is considered one of the main factors for world-wide environmental disturbances ([2]; United Nations Framework Convention on Climate Change 1992). Consequently, natural forests are getting under increased pressure to maintain their ecosystem services (air purification, water purification, soil conservation and stabilisation, animal habitat provision, etc.), protect above and below-ground biodiversity, and maintain productivity (timber, fuel, food source) for livelihoods, and function as net carbon sinks. Healthy trees can act as a local climate buffer and reduce energy costs around farmsteads for example. Urban forests are also gaining in relevance to cool down cities and increase human well-being. Indeed, tree planting projects involving reforestation and afforestation efforts, and the 2 billion tree project over 10 yrs announced in 2021 by the Government of Canada could be mentioned here as an example, are considered as a viable solution to address the effects of climate change through trees’ carbon storage capacity. Yet, these future forests need to be resilient under regimes of future climate (changes in precipitation, temperature rise) and disturbances (pests and pathogens). Physiologically stressed forests would have a reduced carbon storage capacity or even turn into net carbon emitters [3] due to biochemical limitations of carbon uptake under the projected higher temperatures for business-as-usual scenarios [4]. While rising temperatures have been shown to advance the growing season with increased carbon assimilation, ongoing global warming is already reversing the effect due to more frequent drought episodes and heatwaves leading to a diminished sink strength [5].

To keep pace with the current speed of climate change in terms of sustainable forest productivity, effective management of forest genetic resources must be developed. The assisted gene flow approach implementing transfer functions [6] shifts genetic material from different provenances towards their future optimal growth conditions to maintain the current level of productivity. However, with change in average annual temperature and precipitation, there is also increased frequency of biotic and other abiotic stresses that will require establishing higher tolerance not only to drought, but also to an increased risk of frost damage due to rapid dehardening under a warming climate, and attacks from various pathogens. For Canada, boreal conifer forests are especially at risk for a reduction in timber volume, and by 2100, the unaffected wood volumes are projected to be lower than those currently harvested [7]. Therefore, the implementation of resiliency measures to cope with these disturbances is paramount for a sustainable forestry. Aside from crucial intervention measures for forest insect pests and pathogens, tree breeding for resistance against these forest nuisances is urgent to limit their burden on forest ecosystems.

To achieve tolerance to various stressors, Janes and Hamilton [8] emphasized the importance of hybrid zones to select individuals with a favourable combination of features from both parental ancestries. Hybridization enhances broader genetic variation and increases the potential for future adaptation which might be important especially under climate change [9]. Additionally, hybrids are usually more adapted under a broader scale of environmental conditions, while the success of hybridization depends on the phylogenetic distance of the parental species [10, 11]. In some cases, however, there are obvious reproductive barriers due to large phylogenetic distance, causing structural or physiological incompatibility of pollen tubes to establish viable embryos [12]. Therefore, species hybridization is not a universal solution for the generation of resilient genotypes for future environments, and its appropriateness needs to be evaluated on a case-by-case basis.

Spruces are the most important reforestation species in Canada, with hundreds of millions of seedlings planted each year. Interior spruce, a natural hybrid, has a long history of introgression between white spruce (*Picea glauca* (Moench) Voss) and Engelmann spruce (*P. engelmannii* Parry ex Engelm.) as shown by the high levels of recombination events uncovered in hybrid populations [13]. Analyses using genetic markers also revealed asymmetric introgression from white spruce into the local Engelmann spruce species as the initial founder events of the Interior spruce hybrid zone, with eastern British Columbia as the centre of this hybrid zone [13, 14]. Still, the ancestral species maintain their integrity due to environmental selection and limited recent interspecific gene flow (ibidem). In fact, both spruce species occupy distinct ecological niches. While the boreal white spruce has an extremely broad longitudinal range of 111°, the habitat of the mountainous Engelmann spruce is fragmented in western North America covering a much narrower longitudinal range of 23° [13, 15]. Pure species occur below 600 m a.s.l. (*P. glauca*) and above 1,800 m a.s.l. (*P. engelmanii*) according to recent hybrid index assessment using genetic markers [16].

Interior spruce is also one of the economically most important species in western Canada and, due to its hybrid nature, can provide genetic resources needed to cope with climate change [17]. The white pine or spruce shoot weevil (*Pissodes strobi* (Peck); Coleoptera: Curculionidae) is one of the most important biotic threats for Interior spruce [18] and the coastal Sitka spruce (*Picea sitchensis* (Bong.) Carr) species [19] reforestation in western Canada. The weevil is most damaging to young trees [20]. Norway spruce (*P. abies* (L.) H. Karst), which dominates the boreal and subalpine coniferous forests in Europe and has been an important plantation species in eastern North America, is also susceptible to the weevil [21]. The white pine weevil’s life cycle starts with adults overwintering in the litter on the ground. From April to May, adults reemerge and start feeding on the terminal shoot. They favour feeding on the bark (phloem tissue) that is close to the dormant terminal buds. Female weevils lay their eggs into the bark of the terminal shoot, and eggs hatch shortly thereafter (7-10 days). The larvae continue feeding on the leader until July when they reach maturity, and adults emerge after 10-15 days and continue feeding on old and new growth [22]. Thus, terminal growth can be severely affected by weevil feeding and significant losses in productivity due to substantial stem deformations are the anticipated consequence, rendering a spruce plantation meant to provide sawlogs at rotation age worthless.

Adding to this, usually it is the tallest and best growing trees that are preferentially infested by the weevil [22]. Adult weevils can persist over several years and the most important natural control of weevil population numbers is through winter mortalities. However, the ongoing shift to milder winters due to climate change might now create the optimal conditions for maintaining higher weevil population numbers and thus increase the frequency of infestation outbreaks. Therefore, resistance against white pine weevil attacks might have to gain higher importance in breeding objectives for spruces.

While conifers have more or less effective defence mechanisms in place [23, 24], the expression of these defences depends on several factors that is the genetic makeup of the host tree and the local environment where host and pest coexist [25]. For example, it was shown for Sitka spruce that synchrony of weevil activity and budburst phenology of the host is determinant for successful oviposition. Faster budburst phenology of the host may contribute to resistance to the white pine weevil [26]. Environmental cues that indirectly influence host preference include overstory shading (shade-grown trees are less preferred [27]) and fertilizer application (fertilized trees due to their increased bark thickness and leader size enhance their attractiveness for weevil oviposition [28, 29]). One of the best studied direct host defences are the bark resin canals related chemical defences [30] that feature oleoresin blends of a great variety of terpenes, generated thanks to an important functional diversity of terpene synthases found in spruces [31]. However, geography and climate may play an important role in the effectiveness of such resin canal defenses, as shown in trees from the northwestern British Columbian zone of introgressions between Sitka and white spruces [32]. Based on the pure species’ biogeography, O’Neill and colleagues concluded that higher *P. glauca* proportions in hybrids meant increased resistance to the weevil. In general, host tree defences are most effective against local pests, while they weaken under relaxed or absent pest pressure. As the climate warms, these local pest populations may expand their ranges to higher latitudes or elevations, where they may encounter naïve hosts [23], that is hosts that did not coevolve with the pest, since they were never exposed to this threat, and therefore, are less likely to express the appropriate defense response.

The selection of initial plus trees entering the breeding program, and the subsequent selection of parental individuals for advanced generation breeding, has favoured trees with superior growth characteristics. Therefore, screening for resistance of seed sources usually occurs posterior to the detection of a pest or pathogen problem within the breeding program. Therefore, a way to perform mass screening for suitable genotypes is to establish new progeny trials from known seed sources and use artificial infestations to assess genetic control of resistance [33]. These trials can further be used to evaluate how resistance (or tolerance) covaries with growth rate. To do so, the growth rates of tree hosts need to be known before a pest or pathogen occurs to avoid any assessment bias. Long-term breeding programs, such as for the Interior spruce complex *Picea glauca* (Moench) Voss × *P. engelmannii* Parry ex Engelm., exist. It was started in 1973 with a seed collection from 173 ‘plus tree’ Interior spruce individuals growing in natural stands across north central British Columbia (Prince George Selection Unit), where subsequently four progeny test sites involving these open-pollinated (OP) families were established, with subsequent evaluation of the genetic control of juvenile height growth [18, 34]. Furthermore, due to the presence of endemic weevil populations at three test sites, resistance rankings of parents based on retrospective weevil damage assessments were performed over the accumulated sustained damages in each of the 173 OP families [18, 35]. These resistance rankings of parents formed the basis for the selection of parents for the progeny trial at Kalamalka Research station (Vernon, British Columbia) that involved controlled crosses that were generated by mating resistant with resistant, resistant with susceptible, and susceptible with susceptible parents (cross types; [35]). At 5 years of age (in the fall of 1999), this entire Interior spruce trial was exposed to herbivory through an artificially augmented weevil population occurrence. This established the first large white pine weevil trial for spruces in Canada, and screening for weevil attacks took place from 2000 to 2003 [35]. The outcome of this progeny assessment confirmed for the most part the original parent groupings into resistant or susceptible [18]; certain bark characteristics were confirmed as potential indirect screening tools for resistance in spruce families [35]. However, stable positive genetic correlations between resistance and height growth would be necessary for an effective multi-trait improvement program [18, 36, 37]. First assessments of the relationship between height growth (the directly selected trait) and weevil resistance (the indirectly assessed trait by recorded attacks within one OP family progeny test site with 4,330 trees from 139 families) were conducted from the late 1980ies to the early 1990ies in juvenile Interior spruce [20, 36]. The assessment of attacks took into account the total attacks recorded and three individual years of height assessments during this period. Results indicated overall a strong negative genetic relationship (r = -0.61) between growth and weevil attack. As noticed by the authors, tree heights might also reflect early leader loss and damage for more susceptible families [36]. However, this represents an important confounding factor rendering tree height the result of the level of resistance to pest attack rather than an independent trait. Therefore, the establishment of dedicated progeny trials, where artificial weevil infestations are the tools to screen for weevil resistance and assess whether growth assessed before the attack could be related to subsequent attack severity, becomes crucial [33, 35].

In the present study we revisit the entire Interior spruce weevil resistance screening trial encompassing 2,964 individual genotypes (42 families) of varying susceptibility, established in the interior of British Columbia [35]. The same trial was visited previously by the authors [38–40] to investigate a two-by-two factorial spruce progeny made up of families widely segregating for resistance (resistant-by-susceptible cross type) and identify individual candidate genes for resistance to the weevil. These studies also pointed at potential pleiotropic relationships between pre-attack growth and pest resistance, thus our interest in revisiting the weevil trial in more detail. Here, we want to clarify the relationships between resistance to the weevil and, for the first time, pre-attack growth within and across all cross types. We also make direct use of information of parental origin and investigate whether hybrid status in Interior spruce could be a good predictor for weevil attacks. Our results may have important implications for the future management of breeding programs in Interior spruce.

## Material and Methods

### Plant material

The original plant material for this study was established by crossing Interior spruce (*Picea glauca* (Moench) Voss × *Picea engelmannii* Parry ex Engelm.) parents, previously selected from wild stands and then screened for resistance to the white pine weevil *Pissodes strobi* (Peck) (Figure 1A) attacks through an open-pollinated progeny test [18]. The selected parents were then clustered into resistant and susceptible groups and three types of controlled crosses (42 crosses in total) were established [35]: 5 resistant mothers × 6 resistant fathers (16 families belonging to the RxR cross type), 6 resistant mothers × 5 susceptible fathers (20 families; RxS cross type) and 3 susceptible mothers × 3 susceptible fathers (6 families; SxS cross type) (Table 1). These families were planted in 1995 in a randomized design in multiple plots (each plot contained 25 individuals per family) within three blocks (resulting in a total family size of 75 individuals) at the Kalamalka Research Station in Vernon, BC, Canada (Latitude – 50.237222, Longitude – 119.274167) (Figure 1B; for the detailed study site layout based on family plots, see [38]). The plant material was assessed for tree heights in 1998 and 1999 at ages 4 yrs and 5 yrs, respectively (HT4, HT5) (Figure 1B); following artificial augmentation of the local weevil population in October 1999, the annual attack in 2000 (A00) was scored as 0 if there was no attack, as 1 if the attack failed to kill the leader, and as 2 if the attack killed the leader (Figure 1B). Additionally, the number of egg punctures (i.e., those feeding punctures that have egg covering faecal plugs and thus indicate the extent of successful egg laying along the terminal leader [38]) in 2000 (E00) was recorded and scored as five discrete classes ([39], and Figure 1B). The spatial distribution of phenotypes at the study site is displayed in Figure 1B and shows clear patchiness relating to the resistance status of each cross (RxR, RxS, SxS, respectively). The clearest pattern was observed for attack severity where the susceptible crosses exposed excess of attacks while the resistant crosses showed patches of no or low levels of attack (Figure 1B – upper left plot, A00). A similar pattern was observed for the spatial distribution of the egg puncture trait (Figure 1B – upper right plot, E00). This patchiness in the distribution of attack phenotypes was less reflected in heights at both ages. Still, several spruce families emerged as exceptionally high performing (Figure 1B - bottom plots). The class variables annual attack and number of egg punctures were transformed to normal scores following [41]. All information on Interior spruce individuals and the raw data is provided in Supplemental file 1.

**Table 1.**
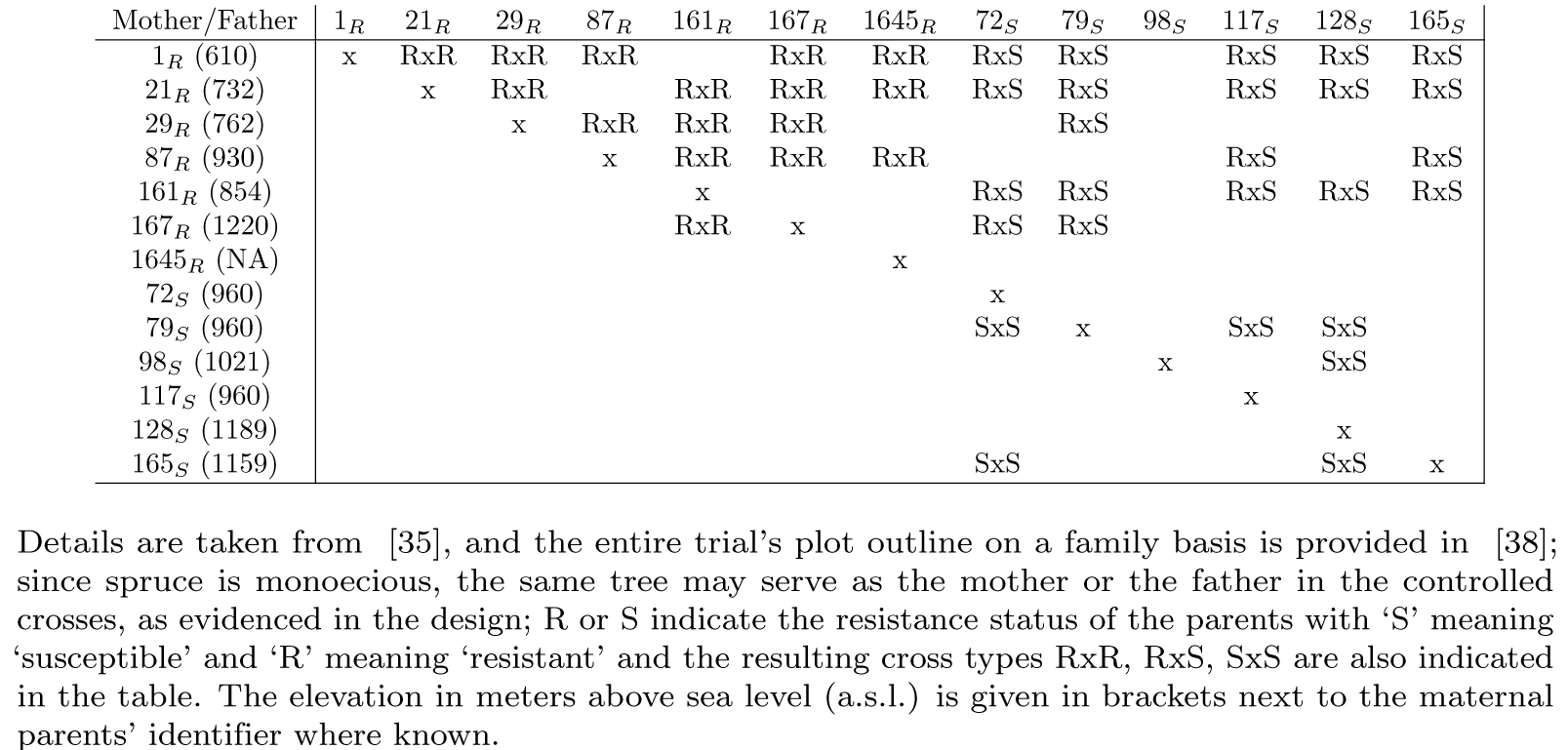
Mating design representing 42 controlled crosses (families) used in this study.

**Fig 1.**
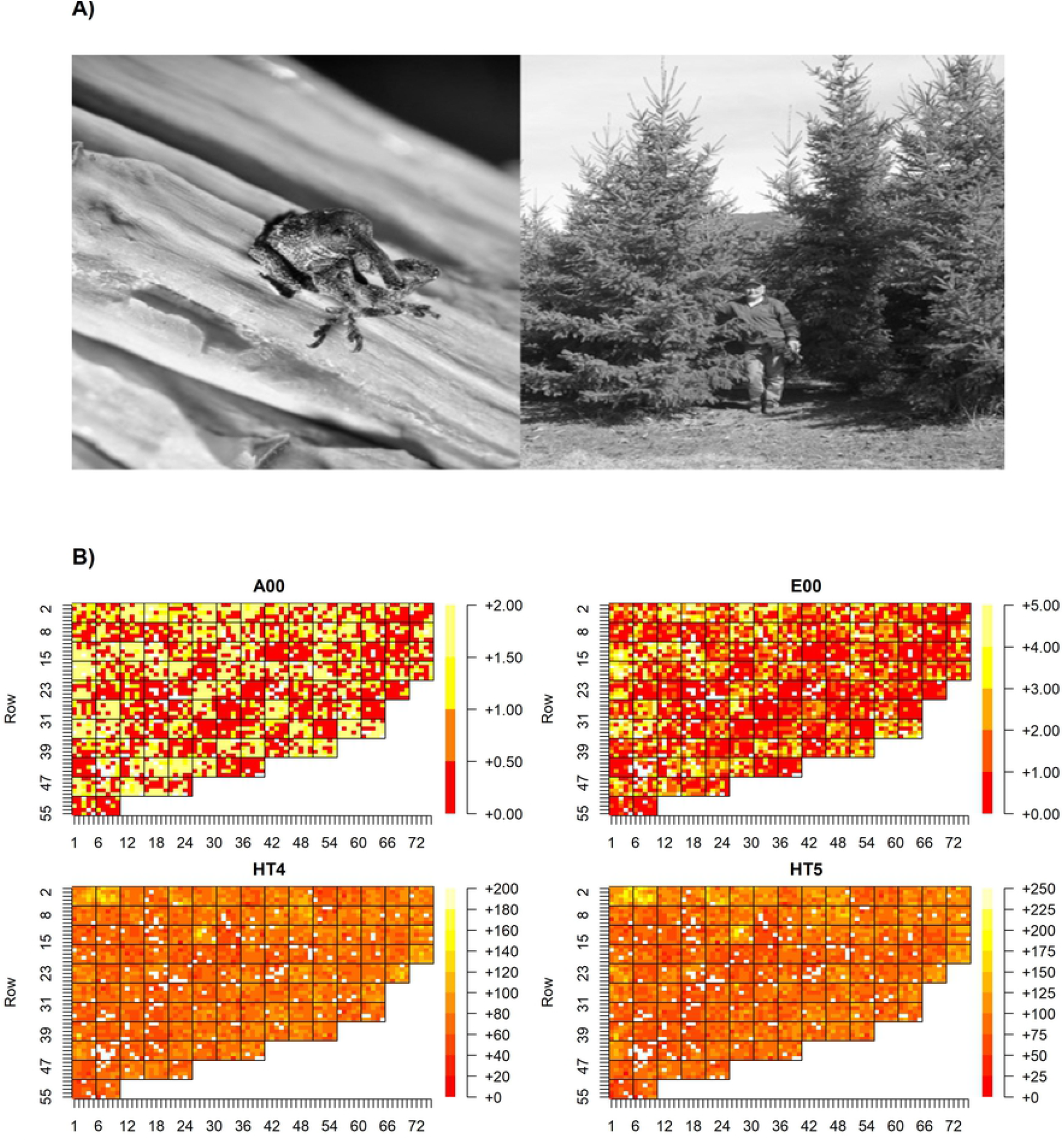
Spruce shoot weevil trial in the interior of British Columbia, Canada. A) Photo to the left: emerging *Pissodes strobi* individual during the weevil rearing experiment (photo credit: Ward Strong); photo to the right: Kalamalka Research Station based Interior spruce weevil trial after thinning (photo credit: Val Ashley).B) Spatial distribution for classes of attack severity (upper left plot, A00), egg punctures occurrence (upper right plot, E00) in the year 2000, as well as for heights measured in cm at four years (bottom left plot, HT4 in the year 1998) and five years of age (bottom right plot, HT5 in the year 1999), respectively, on an individual tree basis within the experimental plot. Family plots are distinguished in the grid, where individual trees are indexed by row and column numbers. The plot outline of the entire trial on a family basis is provided in [38]. Forty-two controlled crosses were generated by mating resistant with resistant, resistant with susceptible, and susceptible with susceptible parents. The ranking in terms of weevil resistance of the individual Interior spruce parents is known and was done previously [18]. The evaluation of attacks was done following artificial weevil infestation of the spruce resistance trial. Height data from pre-attack years are shown. The colour code for ascending phenotypic values (that is for classes of attack severity, egg punctures occurrence, heights) is provided to the right of each plot outline. No formal statistical analysis on the spatial distribution of the weevil attack severity was attempted for this experiment designed as family-based plots where each plot contained 25 individuals of the same family.

### Statistical evaluation

The multivariate mixed linear model using MCMC algorithm implemented in the JWAS package [42] was used to obtain the posterior distribution of variance/covariance parameters as follows:

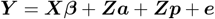

where ***Y*** is a matrix of phenotypes (attack, egg punctures, height at ages 4 yrs and 5 yrs), ***β*** is a vector of fixed effects including intercept and replication effect, ***a*** is the vector of random additive genetic effects following var(***a***) ∼ N(0,G1), where G1 is additive genetic variance-covariance structure following 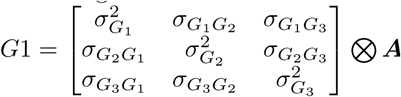, where 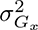 is the additive genetic variance of *x*^*th*^ trait, 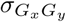 is additive genetic covariance between *x*^*th*^ and *y*^*th*^ trait, ⊗ is Kronecker product and ***A*** is average numerator relationship matrix [43], ***p*** is a vector of random plot effects follo?wing var(***p***)∼N(0,G2), where G2 is plot variance-covariance structure 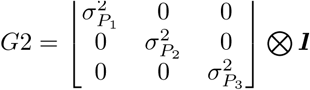, where 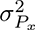 is plot variance of *x*^*th*^ trait and ***I*** is the identity matrix, ***e*** is a vector of random residual effects following var(***e***) ∼ N(0,R), where R is residual variance-covariance structure following 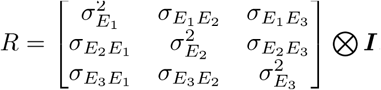, where 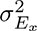 is the residual variance of *x*^*th*^ trait, 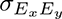 is residual covariance between *x*^*th*^ and *y*^*th*^ trait, ⊗ is Kronecker product, ***X*** and ***Z*** are incidence matrices assigning effects in vectors ***β, a*** and ***p*** to phenotypes of matrix ***Y***. This model was performed for each cross type separately. The model convergence was investigated by the Gelman-Rubin method [44] through merging 5 MCMC runs using 120,000 runs with a burn-in period of 20,000 and thinning of 10 implemented in R package “coda”. The posterior estimates of genetic and environmental correlations between traits were estimated as follows:

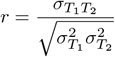

where 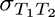 is the covariance between first and second trait, 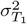 and 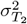 are the variances for the first and the second trait, respectively. The parameters were extracted from the G1 structure for genetic and the R structure for environmental correlations. Phenotypic correlations were estimated by using the original phenotypic data for continuous variables or normal scores for class variables. Since Interior spruce is the natural hybrid of white and Engelmann spruce with an elevational hybridization gradient, we tested the statistical significance of the origin of mother/father trees (in terms of elevation) and cross type on the investigated traits through ordinal logistic regression for class variables (A00; E00) using “polr” function implemented in R package “MASS” as follows:

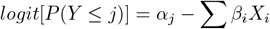

where j is the level of an ordered category (attack severity or egg puncture occurrence) and i is the level of the independent variable (elevation at origin or cross type). Elevation was directly used to order the origin of the parents, while the cross type was recoded into 0 for SxS, 1 for RxS and 2 for RxR. The proportion of total variance explained by the model was estimated in R package “pscl” as pseudo *R*^2^ by comparing the log-likelihood for the fitted model against the log-likelihood of a null model with no predictors implemented. Similarly, the linear regression was performed for continuous variables (HT4; HT5) using the “lm” function implemented in the R package “base” [45]. The proportion of total variance explained by the model was reported as *R*^2^.

## Results

Previous resistance rankings of parental trees indicated that the weevil attack severity in Interior spruce was lowest for the RxR cross type while moderate for the RxS type and strongest in the SxS class (Figure 2A, upper left plot, displayed as the overall percentages for each of the three attack levels). Similarly, the lowest egg puncture occurrence was found in the RxR type while moderate in the RxS type and highest in the SxS type (Figure 2A, upper right plot, displayed as the overall percentages for each of the six egg puncture occurrence classes). Results from ordinal logistic regression showed the statistical significance of cross type on both attack severity and egg puncture occurrence. Additionally, cross type explained 15% and 12% of the total variance in A00 and E00, respectively (Figure 2B). As expected, the predicted effects of cross type for each level of attack severity showed an increase in the likelihood of no attack (A00=0) while a decrease in the likelihood of severe attack (A00=2) with increasing level of cross type resistance (from SxS (recoded as 0) to RxR (recoded as 2)). On the other hand, moderate levels of attack severity (classified as A00=1) did not show any pattern in likelihood related to cross type (Figure S1, Supplemental file 2). For egg puncture occurrences (E00), and with increasing level of cross type resistance, the likelihood for egg puncture level 0 increased while likelihood for all egg puncture levels equal or above 2 decreased. For egg puncture level 1, we did not find any pattern in likelihood related to cross type (Figure S2, Supplemental file 2). Heights at ages 4 yrs and 5 yrs followed a similar pattern (Figure 2A, bottom left and right plots), where families in the RxR class showed the highest median values and which gradually decreased toward SxS class. We noticed that the dispersion of values was highest for RxR class, but lowest for SxS class among all three cross types, potentially reflecting the lowest sample size available for SxS class (Figure 2A). Results from linear regression confirmed the effect of cross type for both height growth years as statistically significant. Yet, the proportion of total variance explained by cross type reached only from 4% (HT5) to 5% (HT4), which was therefore much lower compared to the total variance explained for the resistance traits. The increase in mean height was around 5 cm for HT4 and 6 cm for HT5, respectively, with an increasing level of cross type resistance from SxS to RxR (Figure 2B).

**Fig 2.**
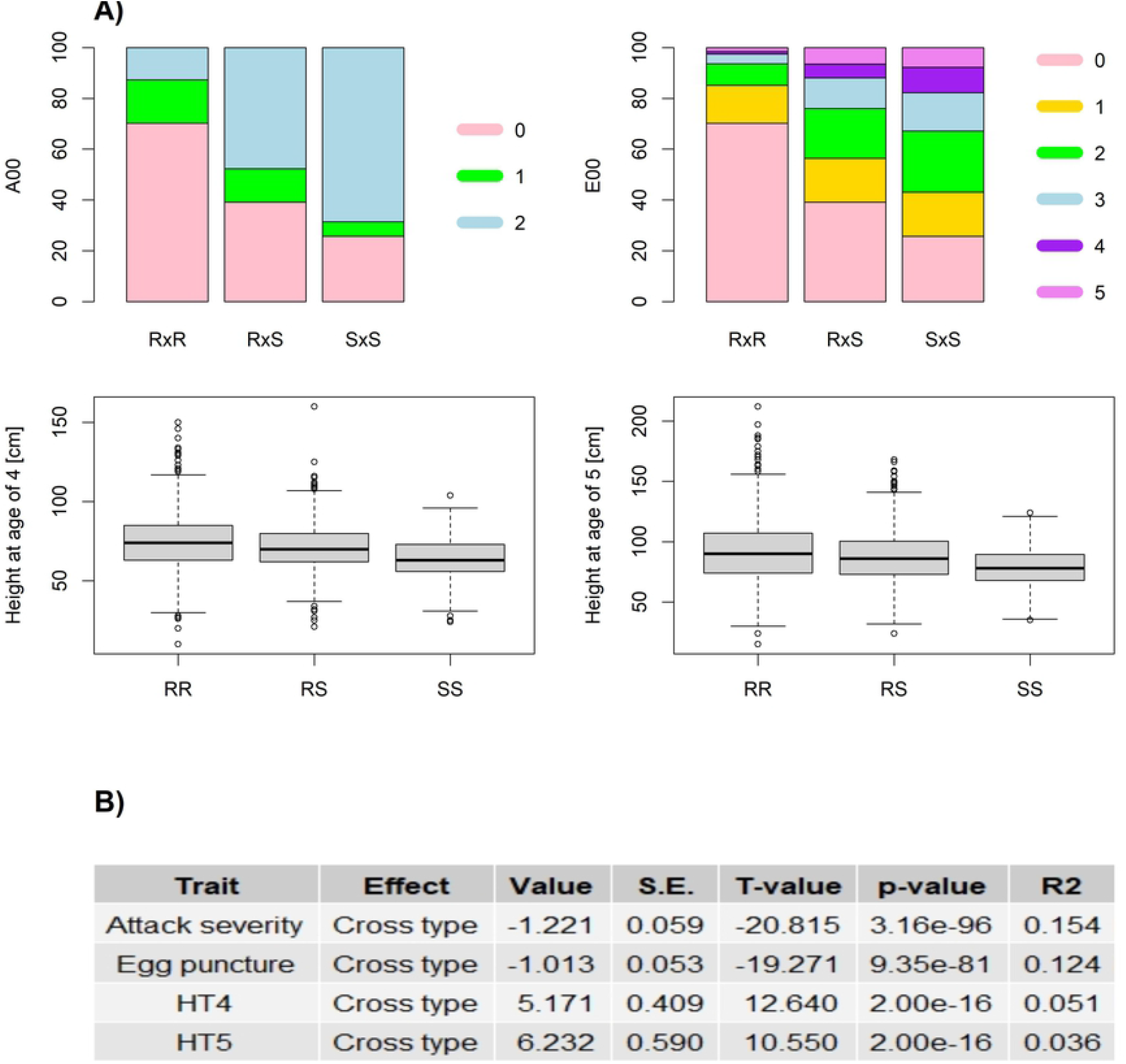
Phenotypic distributions of attack severity, egg puncture occurrences and pre-attack heights within cross types and their significance. A) Stacked bar charts for weevil attack severity (left, upper plot) and egg puncture occurrence (right, upper plot) in Interior spruce following artificial weevil infestation displayed as the overall percentages for each of the indicated severity levels per cross type (RxR, RxS, SxS). The colour code for ascending phenotypic values (that is for classes of attack severity, egg punctures occurrence) is provided to the right of both plots. Furthermore, the boxplots diagrams depicting the dispersion of pre-attack heights at 4 years (left, lower plot) and 5 years of age (right, lower plot) are shown for each cross type of the Interior spruce weevil resistance screening trial. The box in each box plot shows the quartile and the median values of the phenotype dispersion. The whiskers show the range of the phenotypic variation in the individual cross types. Outliers are depicted as points. B) Statistical significance is shown for cross type effect on each investigated trait; HT4, HT5: height at 4 years and 5 years of age, respectively; S.E. represents the standard error and R2 the proportion of total variance explained by the model as pseudo *R*^2^ for ‘Attack severity’ and ‘Egg puncture’ and as classical *R*^2^ for ‘HT4’ and ‘HT5’.

Since Interior spruce genetics reflects the natural hybridization between Engelmann and white spruces along an elevation gradient, we investigated whether the distribution of attack severity in the population can be attributed to parental origins. We found that the mother trees originating from elevations between 600 m and 950 m mostly contributed to only mild attacks, while mothers originating from elevations between 950 m and 1,200 m contributed to most attack severity (Figure 3A, upper row and left column). A similar pattern was observed when the paternal contribution was considered (Figure 3A, upper row and right column). We tested the effect of parental origin on attack severity and found both maternal and paternal effects as statistically significant. Yet, the model explained only 5% of the total variance (Figure 3B). When we plotted the predicted effects of parental origin from ordinal logistic regression, we found that the increase in elevation decreased the likelihood of no attack (A00=0) while increased the likelihood of severe attack (A00=2). No pattern was observed for moderate attack severity (A00=1). These patterns were consistent for both parents (Figure S3 and S4, Supplemental file 2).

**Fig 3.**
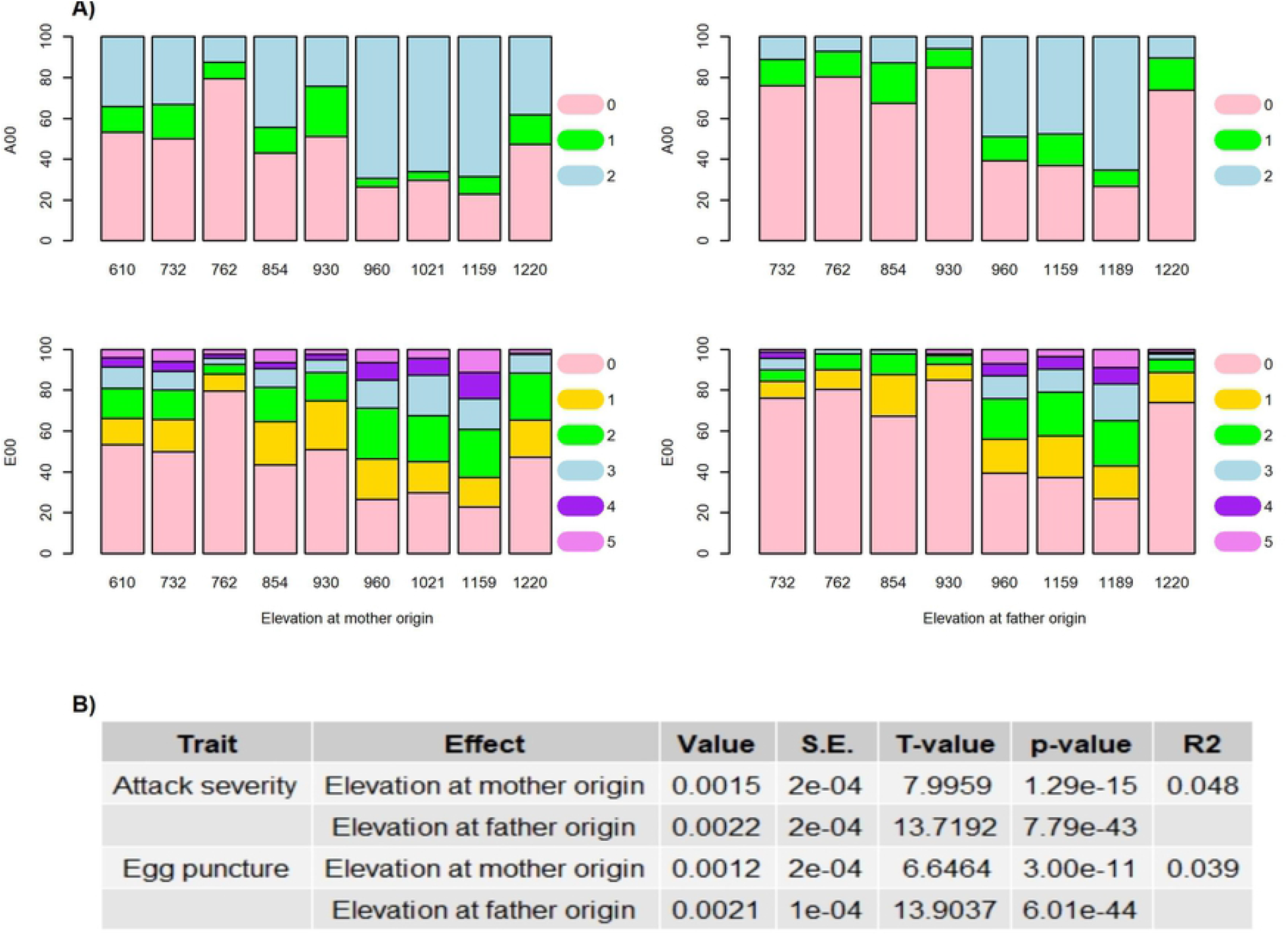
Phenotypic distributions of attack severity and egg puncture occurrences and their significance according to parental origins. A) The distribution of attack severity (A00, upper row) and egg punctures occurrences (E00, lower row) in the Interior spruce progenies (42 families) according to elevation at mother origin (left column) or at father origin (right column) is shown. These 42 controlled crosses were generated by mating resistant with resistant, resistant with susceptible, and susceptible with susceptible parents. The ranking in terms of weevil resistance of the individual Interior spruce parents is known and was done previously [18]. Attack severity increases from 0 to class 2. Egg puncture occurrence increases from 0 to class 5. The evaluation of attack severity and of egg puncture occurrence was done following artificial weevil infestation of the spruce resistance trial. B) Statistical significance of mother/father origin on attack severity and egg puncture occurrence is obtained from ordinal logistic regression. S.E. represents the standard error and *R*^2^ the proportion of total variance explained by the model as pseudo *R*^2^.

For the distribution of egg puncture occurrences in the population according to parental origin (Figure 3A, lower row), we found a less clear switch in the pattern above 950 m with regards to an impact of parent’s origin. This could be explained by a low number of egg punctures that might have already caused damage to the terminal shoots of the offspring, likely due to a lack of resistance mechanisms passed on from parents coming from high elevation (Figure 3A, lower row). The fact that there was only a small proportion of trees with parental origin at 950 m to 1200 m elevation for which egg punctures did not occur, would exactly reflect such pattern. We also noticed for (especially paternal) parents sourced from 1,220 m elevation a drastic drop in weevil attacks (Figure 3A). Again, these patterns were tested via ordinal logistic regression and found to be statistically significant. The model explained 4% of the total variance (Figure 3B). When the predicted effects of parental origin were plotted for each level of E00, we found that the increase in elevation of parental origin decreased the likelihood of no egg puncture occurrence (E00=0), while increased the likelihood at all other levels of E00 (that is E00=1 to 5), although the changes were only marginal. Again, such patterns were consistent for both parents (Figure S5 and S6, Supplemental file 2).

We assessed heritability, genetic, environmental, and phenotypic correlations across and within three distinct cross types (RxR; RxS; SxS) involving nearly 3k genotypes. Results are summarized in Table 2. The narrow-sense heritability estimated across the entire population reached heritability from 0.14 to 0.15 in traits related to herbivory (A00 and E00) and 0.87 related to productivity traits (HT4 and HT5), respectively. When the population was split into cross types, the highest heritabilities were obtained for resistant RxR and susceptible SxS classes ranging from 0.31 to 0.38 in herbivory resistance traits and from 0.86 to 0.88 in growth traits, respectively. However, the SxS class estimates showed larger standard deviations reflecting its smaller sample size (less crosses were available for this particular class). The lowest heritability estimates were obtained for the RxS class reaching only from 0.14 to 0.16 in herbivory resistance traits and from 0.66 to 0.69 in growth traits, respectively.

**Table 2.** Genetic parameters (correlations; genetic control) were estimated for each cross type in Interior spruce.

| Parameter | Trait | All crosses | RxR | RxS | SxS |
| --- | --- | --- | --- | --- | --- |
| Genetic corr. | A00-E00 | 0.60 (0.15) | 0.83 (0.09) | 0.53 (0.18) | 0.62 (0.26) |
| Genetic corr. | HT4-HT5 | 0.96 (0.01) | 0.97 (0.01) | 0.94 (0.02) | 0.96 (0.01) |
| Genetic corr. | A00-HT4 | 0.22 (0.31) | 0.57 (0.18) | -0.54 (0.25) | 0.40 (0.55) |
| Genetic corr. | A00-HT5 | 0.23 (0.30) | 0.56 (0.17) | -0.53 (0.25) | 0.41 (0.54) |
| Genetic corr. | E00-HT4 | 0.33 (0.27) | 0.62 (0.15) | -0.42 (0.29) | 0.49 (0.46) |
| Genetic corr. | E00-HT5 | 0.37 (0.26) | 0.63 (0.14) | -0.38 (0.29) | 0.54 (0.44) |
| Environmental corr. | A00-E00 | 0.92 (0.01) | 0.95 (0.02) | 0.92 (0.01) | 0.88 (0.05) |
| Environmental corr. | HT4-HT5 | 0.75 (0.11) | 0.79 (0.09) | 0.90 (0.04) | 0.76 (0.16) |
| Environmental corr. | A00-HT4 | 0.18 (0.34) | -0.51 (0.23) | 0.65 (0.17) | -0.03 (0.57) |
| Environmental corr. | A00-HT5 | 0.29 (0.32) | -0.48 (0.24) | 0.71 (0.15) | 0.13 (0.56) |
| Environmental corr. | E00-HT4 | 0.19 (0.35) | -0.50 (0.23) | 0.66 (0.17) | -0.05 (0.58) |
| Environmental corr. | E00-HT5 | 0.32 (0.32) | -0.46 (0.24) | 0.73 (0.14) | 0.17 (0.55) |
| Phenotypic corr. | A00-E00 | 0.83 | 0.87 | 0.80 | 0.74 |
| Phenotypic corr. | HT4-HT5 | 0.94 | 0.95 | 0.93 | 0.93 |
| Phenotypic corr. | A00-HT4 | 0.05 | 0.12 | 0.17 | 0.22 |
| Phenotypic corr. | A00-HT5 | 0.12 | 0.16 | 0.24 | 0.31 |
| Phenotypic corr. | E00-HT4 | 0.10 | 0.18 | 0.19 | 0.26 |
| Phenotypic corr. | E00-HT5 | 0.19 | 0.24 | 0.29 | 0.41 |
| $h^2$ | A00 | 0.14 (0.05) | 0.31 (0.14) | 0.16 (0.07) | 0.34 (0.17) |
| $h^2$ | E00 | 0.15 (0.06) | 0.33 (0.14) | 0.14 (0.05) | 0.38 (0.19) |
| $h^2$ | HT4 | 0.87 (0.06) | 0.87 (0.07) | 0.69 (0.14) | 0.86 (0.09) |
| $h^2$ | HT5 | 0.87 (0.06) | 0.88 (0.07) | 0.66 (0.14) | 0.86 (0.11) |

A00: attack severity in year 2000; E00: egg punctures in year 2000; HT4: height at 4 years of age; HT5: height at 5 years of age; $h^2$ : narrow-sense heritability; R...resistant; S...susceptible. Numbers in brackets show the estimated standard errors.

The highest positive genetic correlations were observed between herbivory traits A00-E00 ranging from 0.53 (RxS class) to 0.83 (RxR class) and between growth traits HT4-HT5 ranging from 0.94 (RxS class) to 0.97 (RxR class). The genetic correlations between herbivory resistance traits and growth traits ranged from moderately to strongly positive in RxR and SxS class (reaching from 0.40 to 0.63) to moderately positive correlations in the entire population analysis (reaching from 0.22 to 0.37) to moderately negative in the RxS class (reaching from -0.38 to -0.54). A similar pattern was observed for the environmental correlations with the largest estimates between disease resistance traits ranging from 0.88 to 0.95 and between growth traits ranging from 0.75 to 0.90. The environmental correlations between disease resistance and growth traits ranged from strongly positive in the RxS class (reaching from 0.65 to 0.73) to low in the SxS class (ranging from -0.05 to 0.17) and again low for the entire population analysis (reaching from 0.18 to 0.32) to moderately negative in the RxR class (reaching from -0.46 to -0.51).

The phenotypic correlations were all positive (Table 2), again, highest between growth (0.94) or between herbivory (0.83) traits and with correlations between A00 and height at both ages highest in SxS crosses (0.22 and 0.31, respectively), followed by RxS (0.17 and 0.24, respectively) and then RxR (0.12 and 0.16, respectively). A similar pattern was observed for E00, but generally, correlations reached higher estimates (Table 2). As expected, overall, genetic correlations were stronger than phenotypic correlations. An important outcome of our re-assessment of the white pine weevil trial was that for the RxS cross type (Table 2), the sign flipped from moderately negative to slightly positive for genetic versus phenotypic correlations between herbivory (A00 and E00) and height traits. Of note is that the respective environmental correlations between herbivory (A00 and E00) and height traits in the same cross type (RxS) were strong and positive, whereas they were otherwise either moderate and negative (RxR) or insignificant (SxS).

## Discussion

### Elevation related to the ancestral origins in Interior spruce contributes to the growth and level of resistance against the white pine weevil

The moderate narrow-sense heritability for resistance against the white pine (or spruce shoot) weevil and the relatively high narrow-sense heritability for tree height provides a potential for an effective response to selection for both traits. We stress here that the parental trees deployed in this study were selected solely on the basis of their resistance/susceptibility to white pine weevil attacks and regardless of their origin [18, 35], while our results indicated that susceptible parental trees were predominantly originating from a higher elevation and *vice versa*. The relatively high narrow-sense heritability might result from high genetic diversity captured within the sampled parental trees originating from elevations 610 to 1220 m a.s.l. Since Interior spruce is a natural hybrid between *Picea glauca* and *P. engelmannii* which occupy a transitional elevation zone [14], both species with different growing patterns might contribute to the relatively high narrow-sense heritability. While *P. engelmannii* occupies subalpine environments requiring adaptation for short growing windows and extended winters with deep snow cover, *P. glauca* occupies a boreal environment that competes for light naturally, which selects for relatively fast-growing genotypes. Individuals within the hybrid zone were found to show advantageous features passed on from parental species, such as autumn cold tolerance (*P. engelmannii*) and fast growth (*P. glauca*) [14]. This trend was also observed in this study, where we confirmed that parents of the RxR class that originated from lower elevations, and thus, trees putatively of high *P. glauca* ancestral proportion were tallest (Figure 2). Our hypothesis is based on the observed patterns, while further verification of the population’s actual introgression patterns through genomic markers is needed. We also note that this observation is based solely on pre-attack growth rates. Rossi and Bousquet [46] found that under controlled greenhouse conditions, northern provenances of black spruce showed earlier, and faster bud break compared to southern origins. This result can be seen also as a proxy for the differential local adaptation along an elevation gradient that involves *P. glauca-P. engelmannii* as in the present study, with *P. engelmannii* exhibiting earlier bud burst and budset [14]. Additionally, Wilkinson [47] found a negative genetic correlation between bud break date and early tree height growth. Although De La Torre et al. [14] stated that *P. glauca* does not show growth advantage until it reaches ten years of age, our study found superiority already at 4 years of age in the RxR class, which presumably has the highest *P. glauca* ancestry due to the elevation of origin [16] for the parents involved in RxR crosses (this study). Yet, the differences between cross types explained only ∼ 4 – 5% of the total variance (Figure 2B). While the potential of the different genomic introgression patterns among *P. glauca* × *P. engelmannii* hybrids to explain the differences in the relationship between growth and weevil resistance among individuals has been proposed earlier [40], future research supported by genomic information should be performed to verify the ancestral contribution of the species to individuals in the different cross types and its relationship to the level of resistance/tolerance to white pine weevil attack.

The analysis of attack severity and taking into account the origin of maternal or paternal parents, showed a clear switch from resistant to susceptible genotypes when the elevation of the parental individual’s origin exceeded 950 m a.s.l. (Figure 3). With regards to the co-evolution of the pest with its host, these higher elevations might in fact represent the limit where the short growing window might not allow for completion of the white pine weevil life cycle, and thus, host populations above this elevation are not adapted to this pest. These would thus represent naïve host trees, having never been exposed to the pest. We also found an apparent increase in tree resistance with 1,220 m a.s.l. elevation, thus a pattern which deviated from the general trend. Resistance status was tested in seven crosses, five RxR and two RxS families, which confirmed the resistance segregation pattern expected for a resistant genotype [35]. Since Pinaceae, including spruces, are outcrossing species as evidenced by high levels of heterozygosity in their genomes [48], and the parental genotypes of the F1 generations tested in the trial had directly been sourced from wild stands, the observed resistance pattern above 1,200 m a.s.l. seems valid. We argue that the spring phenology of the hosts might be the determinant factor here. Alfaro et al. [26] found for Sitka spruce that those families resistant to the white pine weevil showed earlier and quicker budburst compared to susceptible families. Generally, individuals at colder sites exhibit greater temperature sensitivity of spring phenology [49]. Therefore, we could assume that families with parental contributions sourced from 1,220 m a.s.l. could show an increased mismatch with the pest’s phenology at the Kalamalka test site, which has a continental climate, and could trigger an accelerated bud flush, since Engelmann spruce and Engelmann spruce-like hybrids require less heat sum [14].

A Québec study performed on 45 Norway spruce genotypes originating from various European provenances, however, found no relationship between budburst phenology, traumatic response intensity and *P. strobi* performance, possibly due to the complete lack of coevolution of this highly sensitive exotic host with the local weevil [50]. The lack of correspondence between the weevil’s biological performance and traumatic resin duct formation in Norway spruce was also suggested in a related study [51]. However, on a broader scale and under a warming climate, the narrowing of phenological mismatches in tree hosts and local insect pests are of concern for the boreal forest [52]. Thus, studies on the state of host-insect phenological synchronies and the forecasted climate-induced phenological changes need to gain in relevance (ibidem). For example, while previous defoliation by herbivores induces earlier budburst in the host as an avoidance strategy [53], warming temperatures again reduce such phenological mismatch due to quicker larval phenology [54].

For our study, we stress here again that the trees were mainly selected for resistance/susceptibility to white pine weevil attacks, and that the observed pattern might also follow clines in adaptive traits. To better understand the role of the introgression level on the resistance/susceptibility status in Interior spruce, genomic information is needed to trace the contribution of the ancestral species. Moreover, to move forward, the implementation of recently available high-throughput assessment methods for tree phenology monitoring that include remote sensing [55] will be crucial to screen trials more holistically and efficiently along trees’ ontogenetic development, thereby offering the possibility to assess phenological changes *in situ*.

### Different resistance mechanisms found between low elevation (RxR) versus low with higher elevation (RxS) crosses

The positive (unfavourable) genetic correlation between tree height and the severity of white pine weevil attack represents a challenge for simultaneous genetic improvement for both traits and some trade-off should be implemented. This is in opposite to cases where tree height is positively correlated with disease resistance such as white pine weevil in Norway spruce [56] or *Leptocybe* gall wasp in *Eucalyptus grandis* [57]. The discrepancy between results from our study compared to those performed in Norway spruce and *Eucalyptus grandis* might be caused by the fact that our sample is at the early ontogenetic stage and has not passed any previous disease outbreaks while the other studies were performed at the advanced ontogenetic stage of the sample. Therefore, the older plant material could pass previous disease outbreaks and the tree height is the result of the level of resistance to pest attack rather than an independent trait. For example, Dungey et al. [58], and later Klápště et al. [59], investigated Swiss needle cast in Douglas-fir introduced in New Zealand. While the Californian provenances were the most productive on South Island which is pest free, their productivity was severely affected by Swiss needle cast on North Island where the Oregon provenances were the most productive due to a higher level of tolerance to the pest.

When each cross type was analyzed separately, opposite patterns in genetic correlations between growth and resistance traits were found. While RxR and SxS classes showed strong or moderate positive correlations, the genetic correlations for the RxS class were always moderately negative. Therefore, fast-growing trees in RxR and SxS were mostly attacked. However, the results of the SxS class should be treated with caution due to the small sample size. The only difference between these two cross types was in the severity of the attack. While the RxR class showed attack severity at level one which indicates that the tree was attacked but that the terminal growth was unaffected, the SxS class resulted in a more severe response (level two) with killed terminal shoots. Surprisingly, the RxS class did not show intermediate results but an opposite relationship between growth and resistance traits. Therefore, fast-growing trees were the least affected by weevil attack in the RxS class. However, the opposite pattern was observed for the environmental compared to the genetic correlations, and these environmental correlations were strong for RxS (with a positive sign) and moderate for RxR (with a negative sign). We argue that a common environment was created by the experimental design where each plot represented multiple individuals that belonged to the same family. This might have generated such rather strong environmental correlations, explained by a strong covariance between the permanent environmental effects in the considered traits [60]. For this reason, phenotypic correlations cannot be used in this case as a surrogate of genetic correlations. Where genetic correlations should have the opposite sign, this is mainly assumed for attributes involved in life history evaluation [61, 62], and synchronous growth is one of the life-history traits in forest trees [63]. Again, also in these cases, phenotypic correlations cannot substitute genetic correlations [60]. While De La Torre et al. [14] already noticed better growth performance and early frost tolerance based on spruce individuals’ hybrid status, our results also indicate that hybrids might indeed be genetically better equipped with an optimized resource allocation between defence and growth. However, our hypothesis should be verified by the re-evaluation of host trees’ productivity at the later ontogenetic stage to better understand the impact of weevil attack severity. Also, the hypothesis about superiority of hybrids has to be further supported by future assessment of the trees’ hybrid index through genetic markers.

As one of the defense mechanisms against bark invading insects, conifer trees are releasing oleoresin around the wound to flood deposited eggs, preventing them from further development into larvae and from causing irreversible terminal shoot damage. A significantly higher density of cortical resin canals was found in resistant compared to susceptible genotypes in Sitka spruce [64] and in white spruce [30]. The same study found a positive relationship between tree growth rate and resin canals density. Porth et al. [40] also found that, overall, genetic correlations between bark histology and growth were all positive. For the RxS class (1,403 genotypes tested), we found indeed those individuals that were fast growing the least attacked. Some families may also display higher tolerance to herbivory, where higher egg puncture occurrence does not automatically result in severe damage outcome for the terminal leader. We found such result, when we regrouped all individual families with strictly 40-80% top-kill outcome [35], where, overall, fast growing trees were less severely attacked and for which genetic correlation between A00 and E00 was extremely low (0.16). However, as stated by Porth et al. [40], the ability to tolerate and at the same time produce well-established chemical defenses towards herbivory might be mutually exclusive, such that we would not expect to observe tolerance in the RxR class.

## Conclusions

Natural disturbances affecting forests are cumulative, highly interdependent and involve biotic stressors, pests, pathogens individually or in association, and environmental factors, drought, fire alike, all of which depend on climate, conditions, which in turn affect phenology or life cycle of these organisms that is trees, pests, pathogens. Climate change along with a warming climate, therefore, presents an increase in the stress on forests due to the expansion of geographic ranges of pests and pathogens to now suitable climates, their increased pathogenicity, enhanced population growth, and potential host jumps [52]. Added to this are increasingly drought stressed and otherwise weakened trees [65]. Therefore, multi-trait selection to build resilience in conifer forests becomes relevant. Moreover, tree planting programs need to increasingly consider the consequences of a warming climate on organisms’ phenologies. The resistance to pests and pathogens plays an essential role in enabling tree adaptation to new ranges through tree population movement. This corroborates that resistance breeding programs are continuously important for managing insect pests and pathogen-related diseases. For example, stable positive genetic correlations between height growth and resistance to a local pest would be necessary for an effective multi-trait tree improvement program. Our study on the widely outplanted Interior spruce concluded that selection of fast-growing hybrid genotypes might be the solution for concurrent improvement in resistance against *P. strobi* attacks, whilst maintaining tree productivity. We hypothesize that the RxS hybrid crosses might have inherited high growth rates and dense resin canals from *P. glauca* but also some other features involved in resistance (or avoidance) mechanisms from *P. engelmannii* preventing severe attack of fast-growing genotypes. Such a feature might be a faster rate of bud flush. This is the first study that clearly highlights that hybrid status might be a good predictor for weevil attacks and that also delineates the boundaries of successful spruce population movement. These results that RxS hybrids might be genetically better equipped with an optimized resource allocation between defence and growth has important implications considering the future management of Interior spruce breeding programs and subsequent spruce plantation programs, which need to ensure that adaptation can be maintained across the environment of deployment. Further work is needed to employ genetic markers in efficient and early tree selection.

## Supporting information

**S1 Fig. Graphical interpretation regarding the ordinal logistic regression for cross type to explain attack severity (A00)**.

**S2 Fig. Graphical interpretation regarding the ordinal logistic regression for cross type to explain egg puncture occurrence (E00)**.

**S3 Fig. Graphical interpretation regarding the ordinal logistic regression for origin of mother to explain attack severity (A00)**.

**S4 Fig. Graphical interpretation regarding the ordinal logistic regression for origin of father to explain attack severity (A00)**.

**S5 Fig. Graphical interpretation regarding the ordinal logistic regression for origin of mother to explain egg puncture occurrence (E00)**.

**S6 Fig. Graphical interpretation regarding the ordinal logistic regression for origin of father to explain egg puncture occurrence (E00)**.

## Acknowledgments

We thank Dr René Alfaro, former, now retired, Canadian Forest Service researcher at the Pacific Forestry Centre, Victoria, BC, Canada, who reared the weevil population, augmented the plantation, and provided egg puncture occurrence data. Gyula Kiss managed the initial parent tree selection, progeny testing, controlled crossing and test site establishment.

## References

1. Weart SR. The discovery of global warming. Harvard University Press 2008.

2. O’Neill BC, Oppenheimer M. Dangerous climate impacts and the Kyoto Protocol. Science 2002;296(5575):1971–1972.

3. Anderson RG, Canadell JG, Randerson JT, Jackson RB, Hungate BA, Baldocchi DD, Ban-Weiss GA, Bonan GB, Caldeira Ken, Cao L. Biophysical considerations in forestry for climate protection. Frontiers in Ecology and the Environment 2011;9(3):174–182.

4. Duffy KA, Schwalm CR, Arcus VL, Koch GW, Liang LL, Schipper LA. How close are we to the temperature tipping point of the terrestrial biosphere?. Science Advances 2021;7(3):eaay1052.

5. Gu H, Qiao Y, Xi Z, Rossi S, Smith NG, Liu J, Chen L. A warmer growing season triggers earlier following spring phenology. bioRxiv 2021;doi: https://doi.org/10.1101/2021.08.08.455549.

6. Aitken SN, Whitlock MC. Assisted gene flow to facilitate local adaptation to climate change. Annual Review of Ecology, Evolution, and Systematics 2013;44:367–388.

7. Aubin I, Boisvert-Marsh L, Kebli H, McKenney D, Pedlar J, Lawrence K, Hogg EH, Boulanger Y, Gauthier S, Ste-Marie C. Tree vulnerability to climate change: improving exposure-based assessments using traits as indicators of sensitivity. Ecosphere 2018;9(2):e02108.

8. Janes JK, Hamilton JA. Mixing it up: the role of hybridization in forest management and conservation under climate change. Forests 2017;8(7):237.

9. Grant PR, Grant BR. Hybridization increases population variation during adaptive radiation. Proceedings of the National Academy of Sciences 2019;116(46):23216–23224.

10. Matute DR, Butler IA, Turissini DA, Coyne JA. A test of the snowball theory for the rate of evolution of hybrid incompatibilities. Science 2010;329(5998):1518–1521.

11. Comeault AA, Matute DR. Genetic divergence and the number of hybridizing species affect the path to homoploid hybrid speciation. Proceedings of the National Academy of Sciences 2018;115(39):9761–9766.

12. Potts BM, Dungey HS. Interspecific hybridization of Eucalyptus: key issues for breeders and geneticists. New Forests 2004;27(2):115–138.

13. De La Torre AR, Roberts DR, Aitken SN. Genome-wide admixture and ecological niche modelling reveal the maintenance of species boundaries despite long history of interspecific gene flow. Molecular Ecology 2014;23(8):2046–2059.

14. De La Torre AR, Wang T, Jaquish B, Aitken SN. Adaptation and exogenous selection in a Picea glauca × Picea engelmannii hybrid zone: implications for forest management under climate change. New Phytologist 2013;201(2):687–699.

15. Degner J. Local Adaptation in the Interior Spruce Hybrid Complex. The Spruce Genome 2020:155–176.

16. De La Torre A, Ingvarsson PK, Aitken SN. Genetic architecture and genomic patterns of gene flow between hybridizing species of Picea. Heredity 2015;115(2):153–164.

17. MacLachlan IR, Yeaman S, Aitken SN. Growth gains from selective breeding in a spruce hybrid zone do not compromise local adaptation to climate. Evolutionary Applications 2018;11(2):166–181.

18. Kiss GK, Yanchuk AD. Preliminary evaluation of genetic variation of weevil resistance in interior spruce in British Columbia. Canadian Journal of Forest Research 1991;21(2):230–234.

19. King JN, Alfaro RI, Cartwright C. Genetic resistance of Sitka spruce (Picea sitchensis) populations to the white pine weevil (Pissodes strobi): distribution of resistance. Forestry 2004;77(4):269–278.

20. Alfaro RI, Fangliang H, Kiss G, King J, Yanchuk A. Resistance of white spruce to white pine weevil: development of a resistance index. Forest Ecology and Management 1996;81(1-3):51–62.

21. Holst MJ. Breeding for weevil resistance in Norway spruce. Z Forstgenetik 1955;4(2):33–37.

22. Alfaro RI. White pine weevil in British Columbia: biology and damage. The white pine weevil: biology, damage and management: Proceedings of symposium, 1994 Jan 19-21; Richmond, B.C., Canada. FRDA Report 226. 1994;7–22.

23. De La Torre AR, Piot A. Liu B, Wilhite B, Weiss M, Porth I. Functional and morphological evolution in gymnosperms: A portrait of implicated gene families. Evolutionary Applications 2020;13(1):210–227.

24. Hammerbacher A, Wright LP, Gershenzon J. Spruce phenolics: biosynthesis and ecological functions. The Spruce Genome 2020:193–214.

25. Ott DS, Davis TS, Mercado JE. Interspecific variation in spruce constitutive and induced defenses in response to a bark beetle–fungal symbiont provides insight into traits associated with resistance. Tree Physiology 2021;41(7):1109–1121.

26. Alfaro RI, Lewis KG, King JN, El-Kassaby YA, Brown G, Smith LD. Budburst phenology of Sitka spruce and its relationship to white pine weevil attack. Forest Ecology and Management 2000;127(1-3):19–29.

27. Taylor SP, Delong C, Alfaro RI, Rankin L. The effects of overstory shading on white pine weevil damage to white spruce and its effects on spruce growth rates. Canadian Journal of Forest Research 1996;26(2)306–312.

28. Manville JF, Sahota TS, Hollmann J, Ibaraki AI. Primary cortex thickness influences the location of ovarian maturation feeding and oviposition of Pissodes strobi (Coleoptera: Curculionidae) within a tree. Environmental Entomology 2002;31(2):198–207.

29. VanAkker L, Alfaro RI, Brockley R. Effects of fertilization on resin canal defences and incidence of Pissodes strobi attack in interior spruce. Canadian Journal of Forest Research 2004;34(4):855–862.

30. Alfaro RI, Fangliang H, Tomlin E, Kiss G. White spruce resistance to white pine weevil related to bark resin canal density. Canadian Journal of Botany 1997;75(4):568–573.

31. Yan X-M, Zhou S-S, Porth IM, Mao J-F. The Terpene Synthase Gene Family in Norway Spruce. The Spruce Genome 2020;177–192.

32. O’Neill GA, Aitken SN, King JN, Alfaro RI. Geographic variation in resin canal defenses in seedlings from the Sitka spruce × white spruce introgression zone. Canadian Journal of Forest Research 2002;32(3):390–400.

33. Alfaro RI, King JN, vanAkker L. Delivering Sitka spruce with resistance against white pine weevil in British Columbia, Canada. Forest Chronicle 2013;89(2):235–245.

34. Kiss G, Yeh FC. Heritability estimates for height for young interior spruce in British Columbia. Canadian Journal of Forest Research 1988;18(2):158–162.

35. Alfaro RI, Jaquish B, King J. Weevil resistance of progeny derived from putatively resistant and susceptible interior spruce parents. Forest Ecology and Management 2004;202(1-3):369–377.

36. King JN, Yanchuk AD, Kiss GK, Alfaro RI. Genetic and phenotypic relationships between weevil (Pissodes strobi) resistance and height growth in spruce populations of British Columbia. Canadian Journal of Forest Research 1997;27(5):732–739.

37. El-Kassaby YA, Ratcliffe B, El-Dien OG, Sun S, Chen C, Cappa EP, Porth IM. Genomic selection in Canadian spruces. The Spruce Genome 2020:115–127.

38. Porth I, Hamberger B, White R, Ritland K. Defense mechanisms against herbivory in Picea: sequence evolution and expression regulation of gene family members in the phenylpropanoid pathway. BMC Genomics 2011;12(1):1–26.

39. Porth I, White R, Jaquish B, Alfaro R, Ritland C, Ritland K. Genetical genomics identifies the genetic architecture for growth and weevil resistance in spruce. PLoS ONE 2012;7(9):e44397.

40. Porth I, White R, Jaquish B, Ritland K. Partial correlation analysis of transcriptomes helps detangle the growth and defense network in spruce. New Phytologist 2018;218(4):1349–1359.

41. Gianola D, Norton HW. Scaling threshold characters. Genetics 1981;99(2):357–364.

42. Cheng H, Fernando R. Garrick D. JWAS: Julia implementation of whole-genome analysis software. Proceedings of the world congress on genetics applied to livestock production 2018;11:859.

43. Wright S. Coefficients of inbreeding and relationship./ The American Naturalist 1922;56(645):330–338.

44. Gelman A, Rubin DB. Inference from iterative simulation using multiple sequences. Statistical Science 1992;7(4):457–472.

45. Team, R Core. R: A language and environment for statistical computing. R foundation for statistical computing. Vienna, Austria. 2021.

46. Rossi S, Bousquet J. The bud break process and its variation among local populations of boreal black spruce. Frontiers in Plant Science 2014;5:574.

47. Wilkinson RC. Inheritance of budbreak and correlation with early height growth in white spruce (Picea glauca) from New England. Forest Service Report NE-391 1977.

48. Mitton JB, Williams CG. Gene flow in conifers. Landscapes, genomics and transgenic conifers 2006:147–168.

49. Prevéy J, Vellend M, Rüger N, Hollister RD, Bjorkman AD, Myers-Smith IH, Elmendorf SC, Clark K, Cooper EJ, Elberling B. Greater temperature sensitivity of plant phenology at colder sites: implications for convergence across northern latitudes. Global Change Biology 2017;23(7):2660–2671.

50. Poulin J, Lavallée R, Mauffette Y, Rioux D. White pine weevil performances in relation to budburst phenology and traumatic resin duct formation in Norway spruce. Agricultural and Forest Entomology 2006;8(2):129–137.

51. Nicole M-C, Zeneli G, Lavallee R, Rioux D, Bauce E, Morency M-J, Fenning TM, Seguin A. White pine weevil (Pissodes strobi) biological performance is unaffected by the jasmonic acid or wound-induced defense response in Norway spruce (Picea abies). Tree Physiology 2006;26(11):1377–1389.

52. Pureswaran DS, D. Grandpré L, Paré D, Taylor A, Barrette M, Morin H, Régniere J, Kneeshaw DD. Climate-induced changes in host tree–insect phenology may drive ecological state-shift in boreal forests. Ecology 2015;96(6):1480–1491.

53. Deslauriers A, Fournier M-P, Cartenì F, Mackay J. Phenological shifts in conifer species stressed by spruce budworm defoliation. Tree Physiology 2019;39(4):590–605.

54. Ren P, Néron V, Rossi S, Liang E, Bouchard M, Deslauriers A. Warming counteracts defoliation-induced mismatch by increasing herbivore-plant phenological synchrony. Global Change Biology 2020;26(4):2072–2080.

55. D’Odorico P, Besik A, Wong CYS, Isabel N, Ensminger I. High-throughput drone-based remote sensing reliably tracks phenology in thousands of conifer seedlings. New Phytologist 2020;226(6):1667–1681.

56. Lenz PRN, Nadeau S, Mottet M-J, Perron M, Isabel N, Beaulieu J, Bousquet J. Multi-trait genomic selection for weevil resistance, growth, and wood quality in Norway spruce. Evolutionary Applications 2020;13(1):76–94.

57. Mphahlele MM, Isik F, Hodge GR, Myburg AA. Genomic breeding for diameter growth and tolerance to Leptocybe gall wasp and Botryosphaeria/Teratosphaeria fungal disease complex in Eucalyptus grandis. Frontiers in Plant Science 2021;12:228.

58. Dungey HS, Low CB, Lee J, Miller MA, Fleet K, Yanchuk AD. Developing breeding and deployment options for Douglas-fir in New Zealand: Breeding for future forest conditions. Silvae Genetica 2012;61(3):104–115.

59. Klápště J, Suontama M, Dungey HS, Telfer EJ, Stovold GT. Modelling of population structure through contemporary groups in genetic evaluation. BMC Genetics 2019;20(1):1–13.

60. Hadfield JD, Nutall A, Osorio D, Owens IPF. Testing the phenotypic gambit: phenotypic, genetic and environmental correlations of colour. Journal of Evolutionary Biology 2007;20(2):549–557.

61. De Jong G, Van Noordwijk AJ. Acquisition and allocation of resources: genetic (co) variances, selection, and life histories. The American Naturalist 1992;139(4):749–770.

62. Lande R. A quantitative genetic theory of life history evolution. Ecology 1982;63(3):6607–615.

63. Lugo AE, Zimmerman JK. Ecological life histories. Tropical tree seed manual. Agricultural Handbook 2002;721:191–213.

64. Tomlin ES, Borden JH. Relationship between leader morphology and resistance or susceptibility of Sitka spruce to the white pine weevil. Canadian Journal of Forest Research 1994;24(4):810–816.

65. Ayres MP, Lombardero MJ. Assessing the consequences of global change for forest disturbance from herbivores and pathogens. Science of the Total Environment 2000;262(3):263–286.

